# TDSTDP

**DOI:** 10.1101/2024.07.21.604454

**Authors:** Bingkun Liu

**Affiliations:** University of Tuebingen, Germany

## Abstract

This is a preliminary thesis on Temporal Difference Spike-Timing Dependent Plasticity (TDSTDP), a variation of STDP that considers dendritic potential. TDSTDP is capable of performing temporal difference (TD) learning without dopamine modulation. Major characteristics of TD learning, including value estimation, value propagation, and temporal shifting of dopamine, are demonstrated in simulations. Additionally, a synaptic calcium model demonstrates its biological plausibility.

## 1 Introduction

The neural implementation of reinforcement learning (RL) in the brain has been a significant research focus, especially after the discovery of temporal difference (TD) learning [29, 32] in the dopaminergic system. It is hypothesized that dopamine (DA) release responds to the prediction error between the predicted and actual reward, while other neural circuits update predictions according to the error signal. Despite the success of TD theory and its descendant theories such as Actor-Critic[33], DDPG[15], and PPO[27], which have been well-developed for extremely complex tasks[36, 3, 31], the investigation of RL neural implementation remains limited to the neural circuitry and synaptic levels.

The learning basis of neural circuits, synaptic plasticity[6], is significantly challenged by the credit assignment problem. The problem, in this context, concerns which synapses should be modified and how they should be adjusted in response to learning signals. The uncertainty arises from two aspects: 1. Temporally, the delayed rewarding signal requires retrospectively identifying neuronal contributors. 2. Spatially, relevant neuronal activities must be distinguished from the coincidental brain activities.

A prevalent solution for both domains is the synaptic eligibility trace in the three-factor learning rule[13, 7, 11, 22, 8]. This mechanism tags recently activated synapses with eligibility traces time-decaying variables that keep track of neuronal interactions in each synapse to determine where and to what extent plasticity should occur. Subsequently, DA acts as a rewarding signal, gating and modulating plasticity based on the eligibility trace.

However, the eligibility trace theory must still endure the biological limitations of DA. DA release, diffusion, and receptor activation enforce a typical latency greater than 100 ms[37, 16, 28], which means any prediction error signal elicited by value estimation must detour back through dopaminergic projections to take effect. Disturbingly, irrelevant neural activity during the detouring period is also tagged. Besides, DA is released in a broadcasting manner[16], i.e. DA diffuses within regions but not at specific synapses. Moreover, the eligibility trace theory struggles to handle negative prediction errors. The assumption employed by [20, 19] is that DA signals below baseline, corresponding to negative error, reverse the polarity of plasticity, whereas such a DA mechanism has not yet been observed (only possible with other modulators like ACh, NE[30]).

In contrast, dendritic potential has negligible latency and uniqueness for individual dendritic branches. It has been proposed to be predictive compartments in various studies[35, 24]. This raises the question: might neurons implement TD learning without DA but with dendritic potential instead?

Here, we propose the theory in which TD learning can be executed through a form of biologically plausible Hebbian-like synaptic plasticity[12], termed TDSTDP. Specifically, TDSTDP, incorporating dendritic potential dependency[10, 25, 35] in Spike-Timing-Dependent Plasticity (STDP[5], a variation of the Hebbian plasticity rule), is capable of calculating TD and propagating values backward through temporally ordered neural circuits.

This theory introduces several novel concepts:

1. TDSTDP, as a local plasticity rule independent of any global signal like DA, guarantees synapse specificity and reduced delay, thereby providing more precise continuous value estimation and updating processes to offer a solution to the credit assignment problem.

2. This theory neither excludes nor contradicts the role of classical DA as a reinforcement signal. Instead, it offers a precise mechanism for DA induction, whereas most other neural learning rule models investigate plasticity in response to the TD signal[13, 8, 7]. Together, they naturally merge into the Actor-Critic model[33], where a network with TDSTDP as Critic estimates value, and the computed TD is conveyed by the DA signal to other brain regions as Actor.

3. An auxiliary biologically detailed NMDAR-Calcium synaptic plasticity model matches TDSTDP. It demonstrates that TD learning can be achieved through simple and biologically plausible synaptic plasticity.

4. The theory capitalizes on the temporal nature of STDP and the robustness of rate-based coding. TDSTDP demonstrates that asymmetric STDP possesses an often-overlooked inherent characteristic of differentiating post-synaptic firing rate changes.

## 2 Fundamental theory

### 2.1 TDSTDP theoretical analysis

The TDSTDP plasticity rule follows the basic principles of Spike-Timing-Dependent Plasticity (STDP): pre-synaptic spikes before a post-synaptic spike (pre-post pairs) lead to Long-Term Potentiation (LTP), while pre-synaptic spikes after a post-synaptic spike (post-pre pairs) lead to Long-Term Depression (LTD). Additionally, LTD is caused by each pre-synaptic spike in proportion to the local dendritic potential, denoted as *ρ*. The diversity of plasticity forms depending on dendritic locations has been observed and modeled in various studies [10, 4, 35, 25], with TDSTDP closely aligning with the findings of Urbanczik et al.[35]. The concern of biological plausibility is addressed by a synaptic calcium model, which will be described in the next section.

Spiking implementation for the following equations is detailed in the Methods Section 6.1. For ease of analysis, TDSTDP is illustrated here from a rate-based perspective. The expectation of weight change over trials can be approximated by the pre/post firing rate within a short period, similar to how classic stimulation-frequency-based plasticity fits into the STDP framework: the post firing rate *post*_*t*+*δ*_ following pre-spikes *pre*_*t*_ results in LTP, while preceding post firing rate *post*_*t−δ*_ results in LTD. The expected synaptic strength *w*_*ji*_ updated by TDSTDP from neuron *i* to *j* can be expressed as:

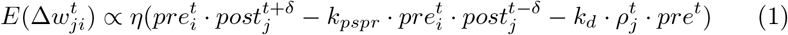

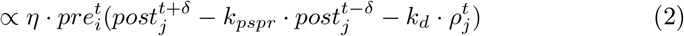

Here, *k*_*pspr*_, *k*_*d*_ are the coefficients for post-pre and dendritic LTD, and *η* is the learning rate. *ρ* is modelled as the integration of local dendrite EPSC, represented by the matrix operation 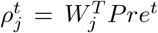 where *W*_*j*_, 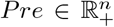 are strength/rate vectors containing all local-dendrite-projecting pre-synaptic neurons.

Considering a postsynaptic neuron *j* is connected by *n* state-encoding neurons (perception, place cell etc, referred to as **state neurons** from now on) with strength *W*_*j*_ at one dendrite branch, as illustrated in Figure 1. Meanwhile, it receives rewarding unconditional stimuli current *I*_*UC*_ and noise through proximal dendrites. Its firing rate *post*_*j*_ can be coarsely approximated by its afferent current (also see Section 6.1):

**Figure 1:**
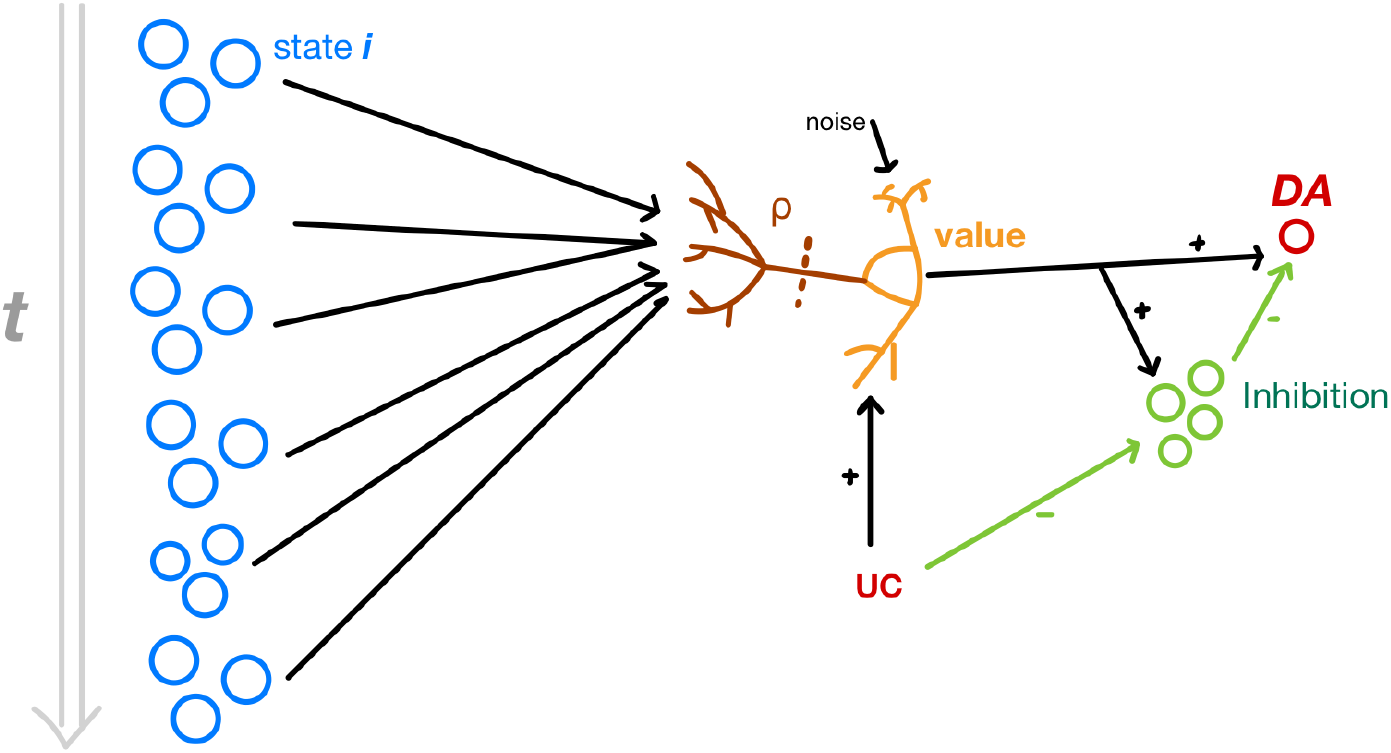
Network model

**Figure 2:**
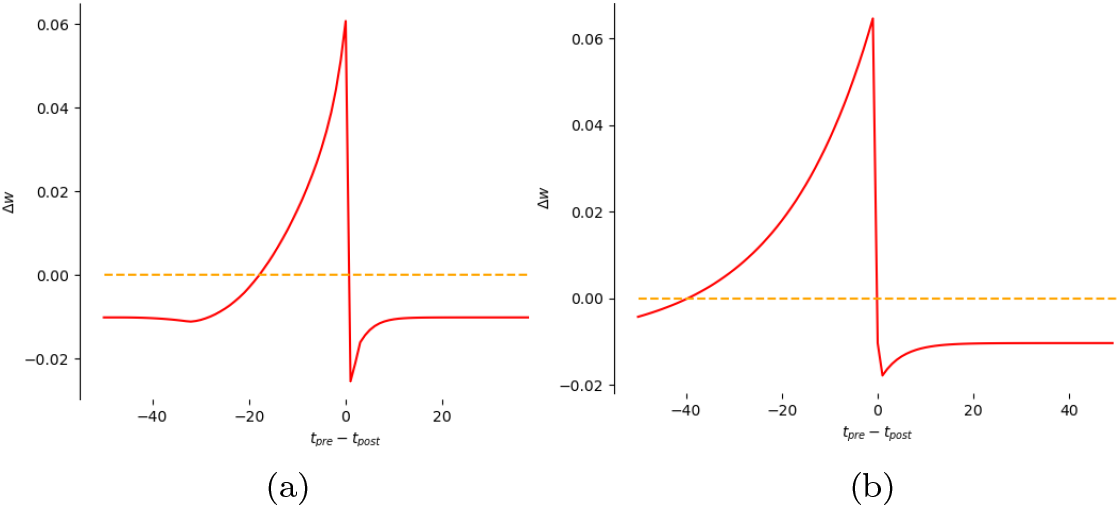
The plasticity modelled by the (a) calcium model greatly conforms with (b) TDSTDP. x-axis is the latency between preand post-synaptic spikes, where negative for pre-post and positive for post-pre. For both simulations, a constant current is injected to model EPSP from neighbour synapses.

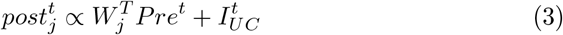

We define 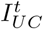 as being proportional to the reward magnitude *r*^*t*^, and the value *V* is estimated by 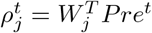: state neuron activation patterns filtered by *W*_*j*_. The post-synaptic neurons, referred to as **value neurons** hereafter, integrate them to efferent an estimator of the sum 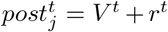, or primarily the value *V* when *r*^*t*^ = 0.

We express *ρ*^*t*^ = *βV* ^*t*^, where *β* is the linear factor converting firing rate to current. Then we continue from equation (2):

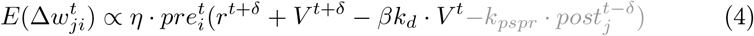

We compare it to the classic TD equation:

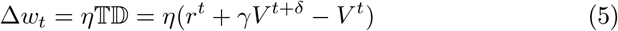

Except for the Post-Pre term in gray, equation (4) is equivalent to TD. Any value decay rate *γ* can be established by an appropriate *k*_*d*_. The Post-Pre term may seem obscure, but it can be excluded by setting *k*_*pspr*_ → 0 to better align with the TD equation. However, a moderate *k*_*pspr*_ not only fits empirical data better but also serves as 𝕋𝔻^*′*^ = *k*_*pspr*_(*post*^*t*+*δ*^ − *post*^*t−δ*^) in a prolonged temporal step size 2*δ*.

Moreover, the scaling factor 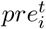 modulates the learning based on presynaptic activity. It ensures synaptic specificity by identifying the contribution from neuron *i*. Mathematically, if each individual state neuron is a continuous feature dimension and states are encoded by feature vectors *Pre*, then 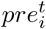 is equiva-lent to the gradient of neuron *i*. Thus, TDSTDP naturally performs gradient descent to minimize 𝕋𝔻^2^:

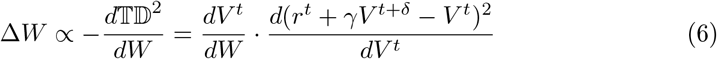

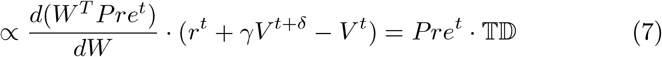

DA release is mediated by a difference-computation neural circuit shown in Figure 1. The value neurons project direct and disynaptic projections with additional latency *δ*_*DA*_ to dopaminergic neurons. The total signal received by dopaminergic neurons is always the difference between time *t* and *t* − *δ*_*DA*_. DA magnitude is also a form of 𝕋𝔻 but provides more stability by smoothing over a longer time scale.

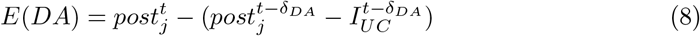

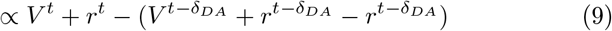

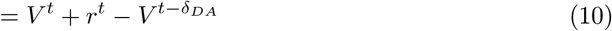

### 2.2 Calcium model

Although TDSTDP differs from STDP measured in conventional in vitro conditions [5], several studies report that the location of synapses on dendrites significantly alters the form of plasticity [25, 4]. Here, We have developed a biologically detailed synaptic plasticity model incorporating NMDAR activation and calcium concentration, which is consistent with TDSTDP.

The calcium model developed here follows basic neurophysiological principles and hypotheses proposed for STDP: pre-synaptic spiking releasing glutamate coincides with raised dendritic membrane potential, unblocking NMDAR receptors to allow calcium influx [21]. Transient back-propagated potential caused by post-synaptic spiking abruptly enhances this process along with intracellular calcium concentration. Moderate calcium concentration leads to LTD, while significantly high calcium concentration leads to LTP [5, 38].

In addition to the original hypothesis, We consider that a dendritic branch can integrate EPSPs with a short time constant, as formulated in Methods. Pre-synaptic spikes lead to subtle depolarization on the local dendritic branch, mostly causing LTD. The occurrence of LTP requires a high calcium concentration threshold that necessitates back-propagated potential to overcome. Therefore, pre-synaptic spikes typically induce LTD, except those closely preceding a post-synaptic spike that induces LTP. A higher afferent current within the local dendrite raises the magnitude of LTD, which is equivalent to *ρ*.

## 3 Result

### 3.1 Linear path simulation

Imagine a mouse moving along a linear path with constant speeds as experiments conducted by Kim et al.[14]. State neurons exhibit Poisson burst firing when the mouse is positioned at specific locations they are mapped to. The burst rates fade in/out with overlap to assemble continuous transportation (fig 3(a)). The unconditional rewarding stimuli (UC) appear when entering one or a few locations leading to transient raising *I*_*UC*_. An example spiking pattern and afferent current are shown in Figure 3(b). State 0 is defined as arbitrary off-path states to mitigate boundary artefact as explained in the stochastic simulation. It is positioned at both the head and tail of the sequence, and its weight is fixed. It can be regarded as the ordinary state when no rewarding relevant cue has been encountered.

**Figure 3:**
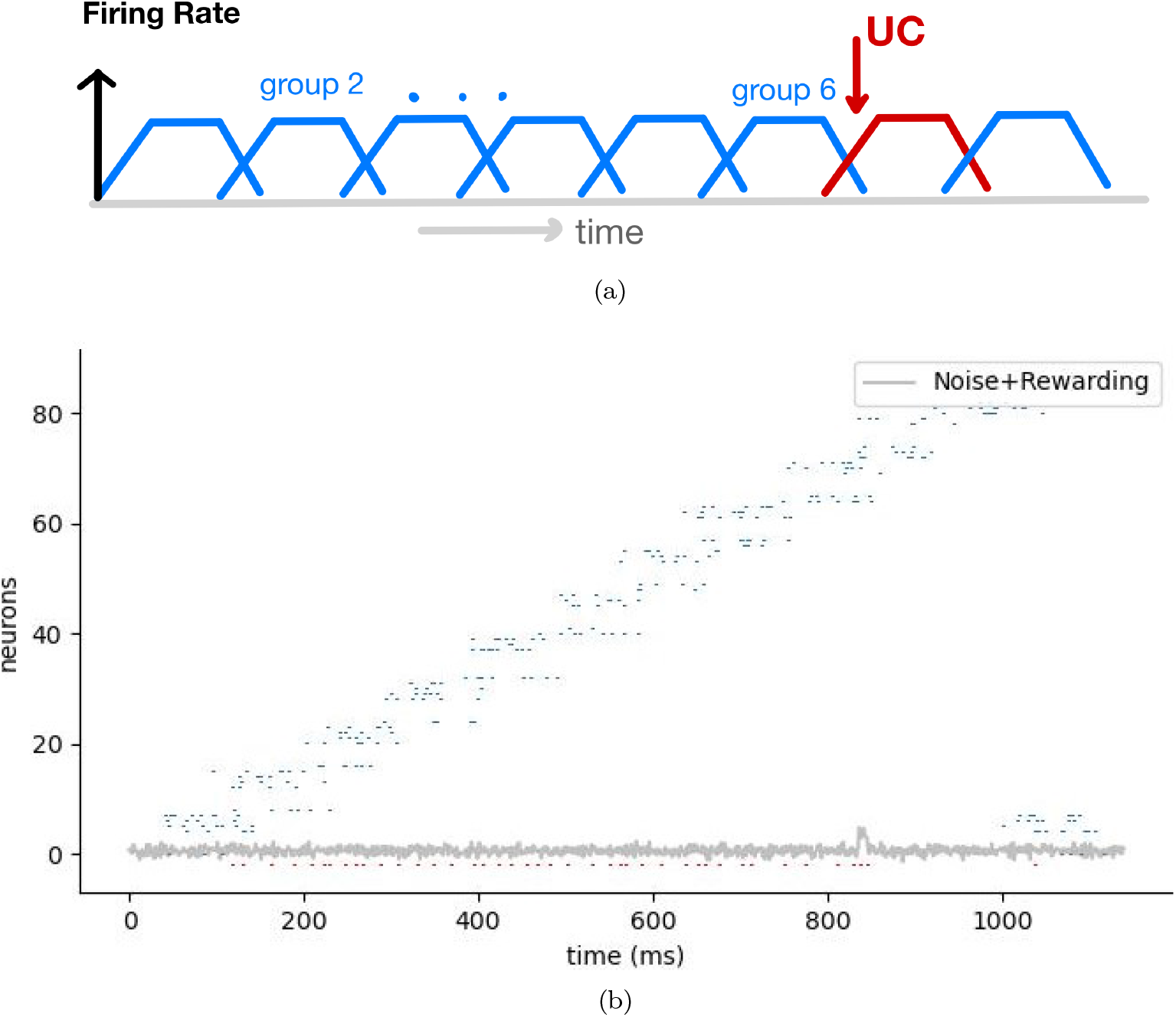
(a) Sequential firing pattern for state neurons along the linear path. The firing rate gradually rises and declines for entering and exiting locations. (b) An example of a spiking pattern and the current variation of the scenario that a reward is given at a certain state, as described in Section 3.1.1. Resulting value neuron spikes are coloured in red.

We simulate TDSTDP with 4 rewarding scenarios to validate the characteristics owned by the original TD theory. TDSTDP is supposed to be capable of

1. propagating value backwards along states but not forwards and shifting DA release timing accordingly,
2. converging to the expectation for stochastic rewards,
3. summing up multiple rewards coming sequentially,
4. decreasing value estimation for reduced or missed reward.

Note that this simulation can be generalized to spatially or temporally ordered states in one dimension.

#### 3.1.1 Plain TD

We start with a scenario where a reward always appears at a certain location during training trials. The simulation results indicate that TDSTDP exhibits a most fundamental TD characteristic: value propagation. As illustrated in Figure 4(b), the value increases for states nearing UC and early states in subsequent trials.

**Figure 4:**
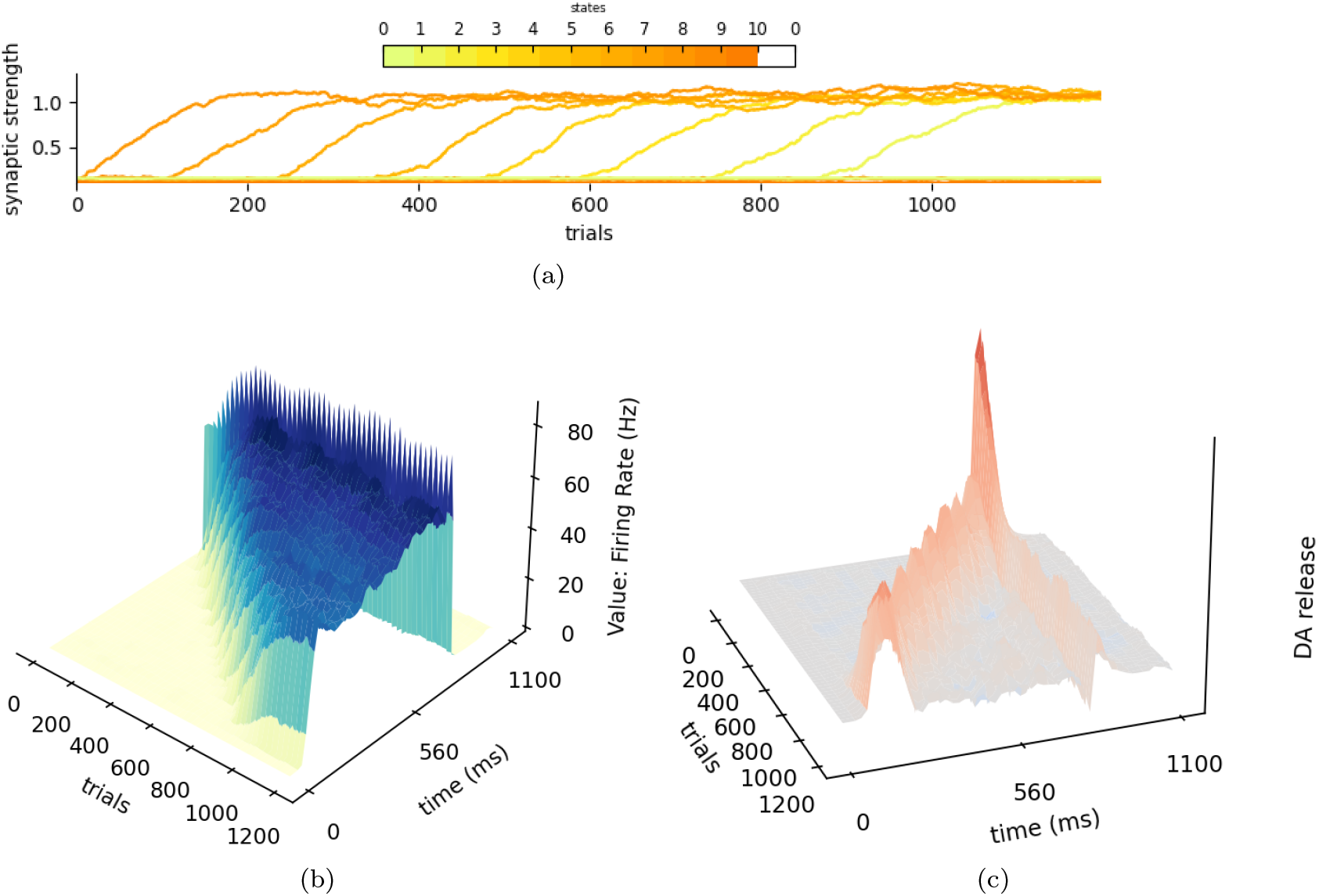
Simulation result for the plain rewarding scenario. (a) State neurons preceding UC increase the synaptic strength in the order of proximity. Deeper colours represent later states. (b) Value neuron firing rate, as the proxy for value, back-propagated in time through trials. (c) DA release conforms to the temporal shift property.

The rise in value is due to the enhancing synaptic strength of the state neu-rons, ordered from those closest to UC (darker) to earlier states (lighter). These values eventually converge to a similar level, indicating the network can learn a stable representation, as analyzed in Section 6.2. Meanwhile, neurons activated after UC decrease their synaptic strength to baseline, ensuring unidirectional value propagation.

The gradual temporal shift of DA release is posited by the original TD theory [29] and recently verified by empirical data [2]. Dopaminergic neuron activity, modelled to represent DA release, is visualized in Figure 4(c). Peaks significantly higher than the baseline are highlighted in red, forming a continuous trajectory from UC to the beginning across trials. This gradual shift of DA release peak strongly proves TDSTDP is consistent with TD theory.

In our results, part of the DA release persists at the time of UC arrival, visible as the vertical lighter red path in the data. This indicates that there is a component of DA release associated with the actual UC arrival rather than TD, thus it does not shift accordingly. This observation deviates from plain TD theory. However, phasic DA increments coinciding with the actual arrival of UC after training have also been extensively documented [14, 37, 2]. Therefore, this phenomenon is supported by empirical evidence and suggests a nuanced neural implementation.

#### 3.1.2 Omission

Omission is a classic demonstration of TD theory. Once the subject has learned to predict the reward, the reward is suddenly reduced or cancelled at the original state. This unexpected reward reduction causes a negative 𝕋𝔻, eliciting DA release below baseline. This negative 𝕋𝔻 should also be back-propagated to adjust the value estimated by preceding states.

To validate the TDSTDP capability of calibrating both magnitude and occurrence of rewards, UC is first degraded by half at trial 1500 and then fully deprived.

Similar to the initial positive learning, the decrement in synaptic strength and value also begins from states just before UC to earlier states. The issue arises because the firing rate of value neurons is not perfectly proportional to the value due to thresholds and other nonlinear factors between current and spikes. Varying reward magnitudes across simulations indicate that the value within a certain range can be well linearly decoded. Empirical evidence [14] indicates imperfect linear relationships between DA release and reward, but this observation does not undermine TDSTDP in any way.

DA release demonstrated in Figure 5(c) produces two negative TD signals (highlighted in blue) instructing preceding states to learn from two reward reductions. This is attributed to concentrations below a baseline level, as DA concentration is a non-negative variable.

**Figure 5:**
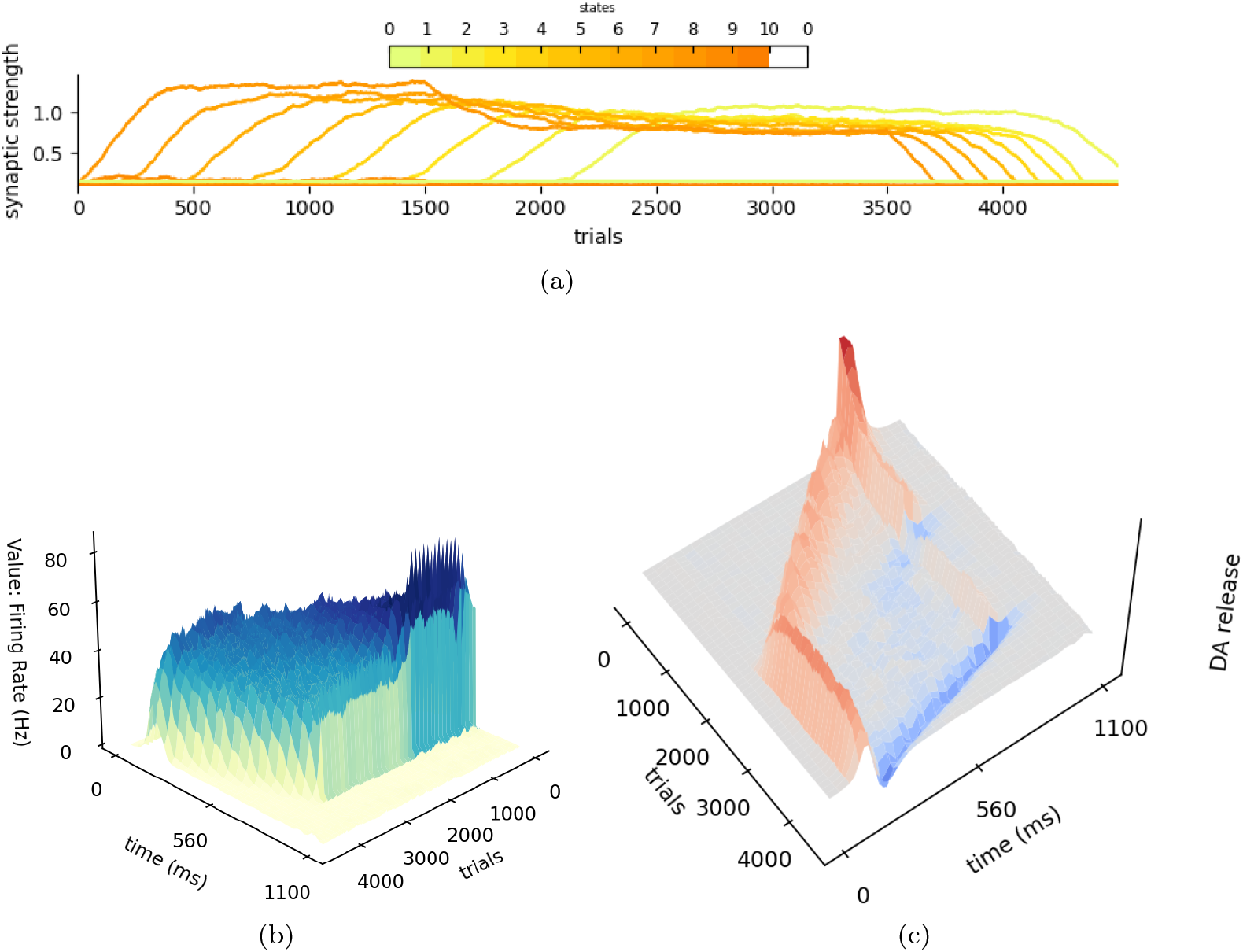
Simulation result for the reward omission scenario. (a) After reward reduction, state neurons preceding UC decrease the synaptic strength in the order of proximity. (b) Value neuron firing rate shows declining slopes and stabilized platforms after two reductions. (c) DA release below baseline temporally shifts backwards. Two reductions produce two negative paths in blue.

#### 3.1.3 Summation of multiple rewards

Suppose there are two rewards, UC1 and UC2, given at two displaced locations. The value when entering the path should anticipate the occurrence of both rewards, though it is not necessary to be the exact summation due to value decay. After receiving UC1, the value should step down to anticipate UC2 alone.

The terrace pattern of value neuron firing rate in Figure 6(b) corroborates our hypothesis. UC1/2 are designed to sum up to UC in plain simulation so that the total amount of rewards is identical for a fair comparison. It can be observed that values at the beginning are comparable, indicating successful summation. Two-reward simulation has a slightly lower value at the beginning because the time displacement is prolonged till UC2.

**Figure 6:**
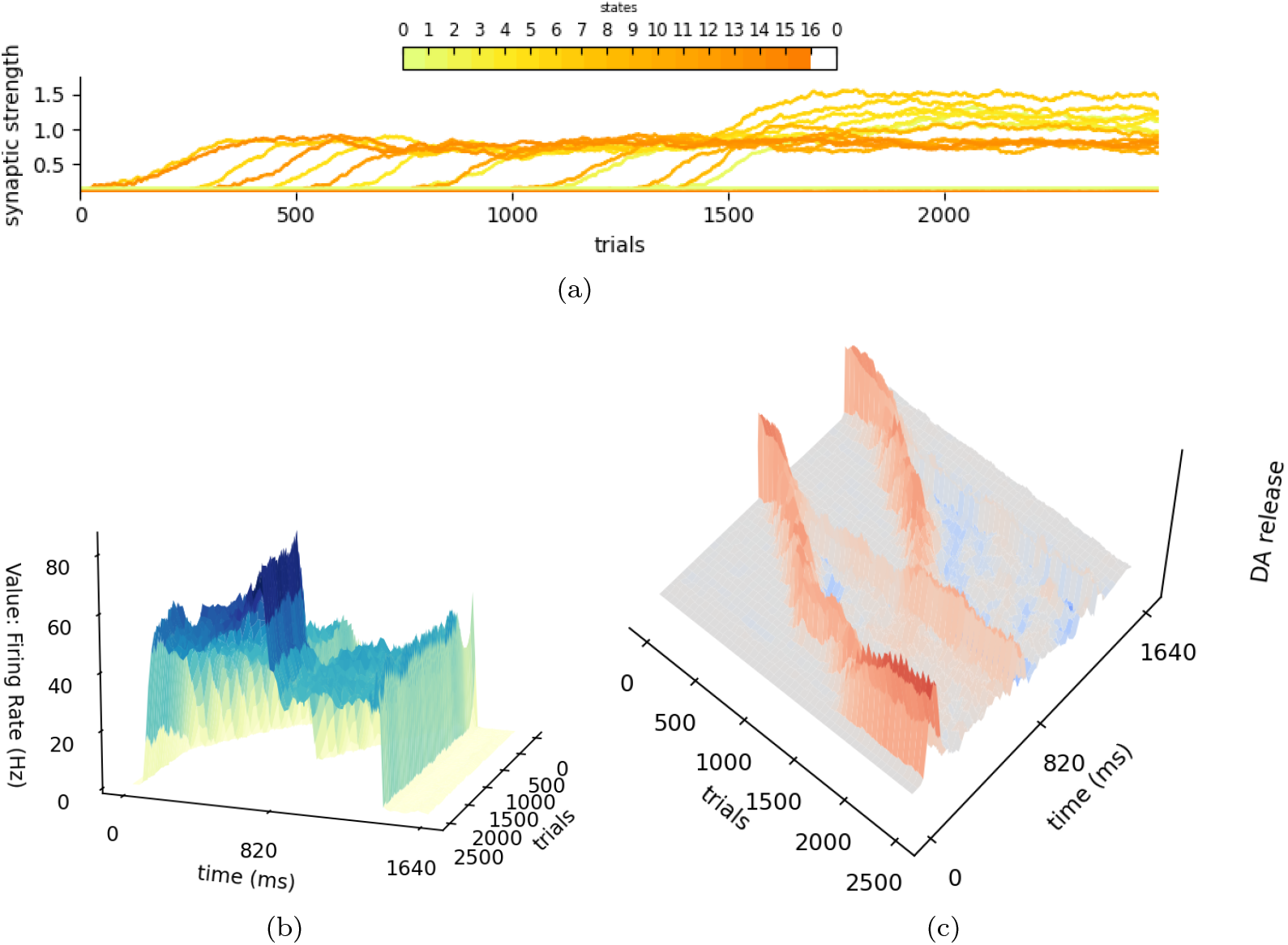
Simulation result for the two-reward scenario. (a)State neurons before UC 1 converge their synaptic strength to a level higher than states between UC 1 and 2. (b) Output Value steps down after UC 1 from reward summation to only UC 2 and steps down to 0 after UC 2. (c) DA temporal shift paths start from two rewarding states separately and merge at the beginning state. The path from UC 2 is sparse after passing UC 1.

**Figure 7:**
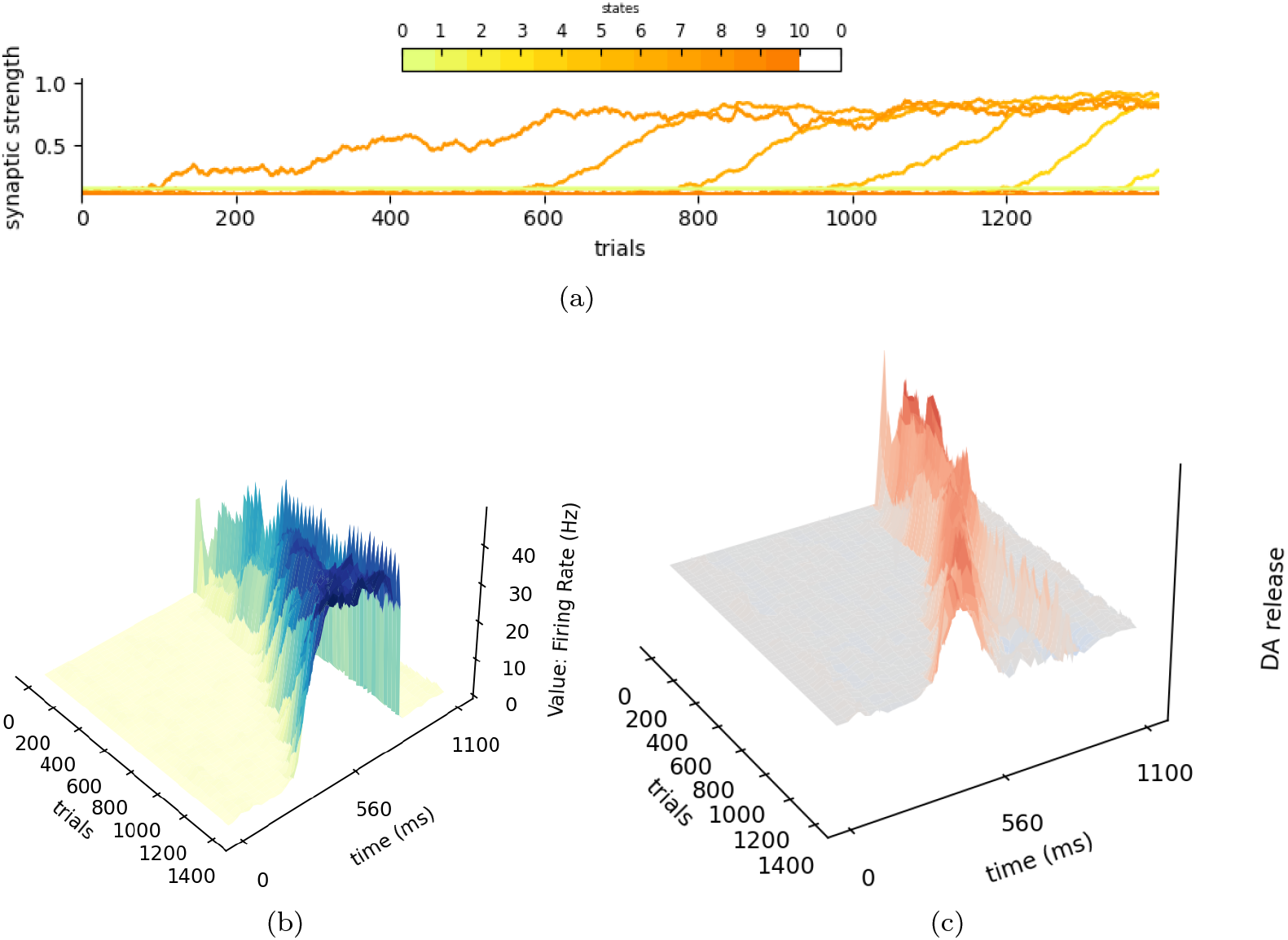
Simulation result for the stochastic reward scenario. (a) The synaptic strength (b) Output Value steps down after UC 1 from reward summation to only UC2 and steps down to 0 after UC 2. (c) DA temporal shift paths start from two rewarding states separately and merge at the beginning state. The path from UC 2 is sparse after passing UC 1.

Consistent with TD theory, DA shift paths originate from each reward and merge at the path entering. Two red trajectories can be identified in Figure 6(c), though the trajectory from UC2 is distorted after overcoming UC1. The distortion is caused by the firing rate non-linearity and UC arrival signalling response as discussed previously.

This observation suggests that TDSTDP can effectively integrate multiple reward signals over displaced locations and temporal delays. The ability to sum and propagate values backwards while maintaining distinct paths for each reward supports the robustness of TDSTDP in handling complex reward structures.

#### 3.1.4 Robustness to Stochastic

When rewards become less deterministic, a valid TD model must robustly converge to expectations. DA has uniform responses between reward probability and magnitude as long as the expected value is identical[34]. TDSTDP can account for stochastic rewards appearing with a probability *p* (*p* = 0.5 in the simulation). The firing rate of value neurons is lower than that under steady presentation of rewards in the first simulation.

Similar to the majority of RL methods, a smaller learning rate *η* helps stabilize learning. For a fair comparison, the same *η* is used in the simulation shown in Figure 4. We observe the propagation of value fluctuations. Furthermore, the speed of propagation and DA shift are reduced even with the same learning rate.

State 0, as defined earlier, represents arbitrary states unrelated to rewards. There are countless such states, each of which has a low probability of being followed by states on the path and is more likely to be followed by another similar state. The expectation of future value diminishes to zero, the same as state 0 is enforced.

It should be noted that TDSTDP is robust to stochastic variations in reward occurrence, timing, and external noise. In all simulations, the exact timing of rewards is uniformly distributed within a 20ms interval. Additionally, value neurons are affected by uncorrelated Gaussian noise *I*_*n*_ (equation 11). We can conclude that TDSTDP is robust to stochastic variations in reward occurrence, timing, and external noise.

## 4 Discussion

### 4.1 Two TD manifestation

One might already notice that TD appears twice in the entire TDSTDP theory: explicitly as DA release and implicitly in the plasticity rule. DA release is produced with the particular network architecture depicted in Figure 1. The inhibitory interneurons cause a time displacement and reverse the sign of the delayed value signal, thereby explicitly establishing the subtraction in TD (equation 5). TD manifested by DA is enforced with the long timescale of this particular architecture and is therefore smoothed by a low-pass filter.

A prerequisite for this architecture is the correct estimation of value, and consequently, the reliability of the TDSTDP rule. Although TDSTDP does not include a TD term in its equation, it is shown to embed TD by deriving its expectation of net plasticity (equation 4). In contrast to DA, TD manifested in the plasticity rule shares the short timescale with STDP. It is subject to stochastic and rapidly varying fluctuations, thus necessitating averaging across trials for stable TD expression.

This manifestation has an advantage over DA in that it does not concern itself with negative TD. Although DA levels below baseline can be an explanation, the dynamic range between 0 and baseline remains constrained. Negative TD expressed as LTD enjoys the same dynamic range as positive TD expressed as LTP.

The two manifestations of TD align with the Actor-Critic model, where TD guides the Actor and Critic differently[33]. Value neurons serve as Critic, instructed by implicit TD. DA released from the network projects to other brain regions, serving as the Actor to instruct policies.

### 4.2 Markov states encode Cue and Timing

Simulations in Section 3.1 assume a task context employed in [14] where stimuli continuously inform the distance to the reward. TD is also frequently examined in delayed reward schematics [29], where rewards appear after a perceptual cue with a certain time lag. In such cases, there is no reward-relevant stimulus within the lag period. This can undoubtedly disrupt TDSTDP, as if state neurons are firing randomly during the lag, no value can be propagated back.

However, as stated, state neurons at this brain level are presumed to encode Markov states. According to Markov chain theory, the value depends solely on the current state, which, in some sense, encompasses all necessary historical information. Exemplified by the state neurons, they should encompass the timing and content of the cues. This mechanism is well explained by recurrent excitation forming working memory [18], though it is beyond the scope of this paper.

Therefore, TDSTDP and the simulation can be generalized to delayed reward scenarios and any other contexts compatible with Markov chain theory. This also partly addresses the distal credit assignment problem, as previous cues are encompassed in the state, weighted by their degree of salience and novelty.

### 4.3 Dendrite as a predictive compartment

Pyramidal neurons typically have apical dendrites that extend far from the soma. This morphological structure segregates intracellular ionic, molecular, and membrane potential transmission[10, 17]. The heterogeneous physiology within a single neuron enables computational functions comparable to multilayer neural networks[1, 17]. In particular, information flows bi-directionally: feed-forward via somatic integration and feedback via back-propagated potential. Richards et al.[25] suggest that dendrites can help solve the credit assignment problem by isolating feed-forward and feedback signals.

Synaptic plasticity, critically modulated by physiological environments, hinges tightly on local dendrite properties[10, 4]. Rao et al.[24] explicitly formulate TD in dendritic plasticity, reproducing an STDP-like rule. Urbanczik et al. [35] propose that a rate-based plasticity rule conforming to TDSTDP allows dendritic potential to predict somatic spiking in a two-compartment model. They demonstrated that the dendrite, as a separate compartment, can be predictive. TDSTDP theory develops this into spiking and even more biologically detailed models, exploiting this characteristic with this particular neural connectome and sequential afferent. Furthermore, it demonstrates that dendritic computation in vivo can perform exact TD learning.

### 4.4 TDSTDP generalize to predictive coding

TD theory shares significant similarity with predictive coding theory (PCT)[23]. The essence of TD is to estimate the value as a prediction of upcoming rewards, which can be aligned with predictive coding for rewards. Meanwhile, prediction error is employed in both theories, with reward prediction error (RPE) manifested as DA release in TD, and predictive error regarded as salient information to be processed in PCT.

Under the same philosophy, TDSTDP as a learning theory correcting postsynaptic firing rate towards future activity can be competent as a learning basis for both TD and PCT. The divergence between the two theories is that TD theory predicts value, an internally generated measure, for most unrewarded moments. PCT, on the other hand, always predicts external stimuli or their representations at various levels. TDSTDP can be adapted to PCT by receiving predictive error through *I*_*UC*_, acting as a regular guiding signal.

### 4.5 TDSTDP bridges rate and temporal coding

The debate between neural rate and temporal coding has brought attention to STDP. Temporal coding posits that information is encoded in precise spike timing rather than firing rates, which require averaging over time. STDP, emphasizing the timing of preand post-synaptic spikes, serves as a crucial mechanism supporting temporal coding. Especially in asymmetric STDP, post-pre pairs lead to plasticity polarity opposite to pre-post pairs. Consequently, most STDP studies focus on isolated spike pairs or triplets.

However, it is often overlooked that each pre-synaptic spike participates in both post-pre and pre-post schemes, serving as a time boundary. The net plasticity is determined by the difference in post-synaptic firing rates, calculated by subtracting the number of spikes before the boundary from those after it. In the context of rate-encoding neural activity, asymmetric STDP effectively learns the derivative of the post-synaptic firing rate in expectation. This functionality is influenced by the causal correlation between preand post-synaptic spikes, which can be mitigated by uncorrelated currents and stochastic firing timings. Nonetheless, this provides promising insights into STDP’s role in noisy rate coding neural circuits.

## 5 Conclusion

This part is left unfinished. Pending for more experiments and content.

## 6 Methods

### 6.1 TDSTDP Spiking formulation

The post-synaptic neuron *j* somatic voltage *U*_*j*_ follows the Leaky-Integrate-Firing dynamic. It has rest membrane potential *θ*_*rest*_ and receives Gaussian noise *I*_*n*_ ∼ *𝒩* (*µ, σ*^2^) :

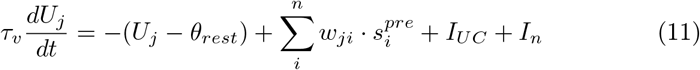

Similar to conventional STDP formulation in spiking simulation, the weight update in TDSTDP is computed as:

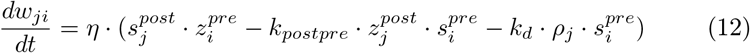

Dendrite potential *ρ* (omit offset *θ*_*rest*_) collecting afferent projecting to the local branch has a short time constant i.e. *τ*_*ρ*_ = 4*ms*:

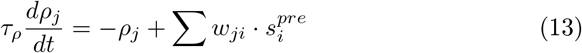

Spike trace *z* records the temporal proximity of previous spikes:

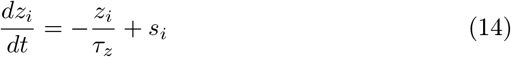

Here *s* = 0*/*1 indicates spiking. *τ* is time constant, *η* is learning rate, and *k*_*pspr/d*_ are constant coefficient constrained by the equilibrium below.

DA concentration has a large time constant *τ*_*DA*_ integrating the difference of *V* + *r* between concurrent and *δ*_*DA*_ ago by delayed inhibition.

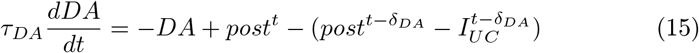

### 6.2 TDSTDP equilibrium

Formulate Equilibrium states to determine *k*_1_ and *k*_2_:

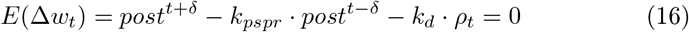

According to value discount, unrewarding steady states have *V* ^*t*^ = *γV* ^*t*+*δ*^:

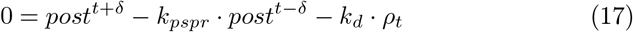

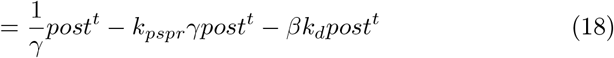

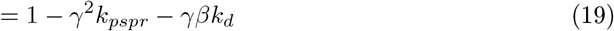

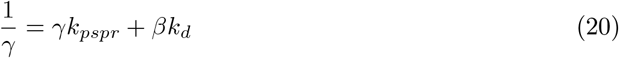

So we have the constraint for setting *k*_*pspr*_, *k*_*d*_. We can verify this at the state before Rewarding:

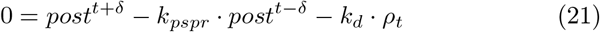

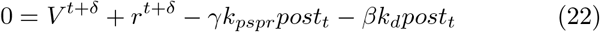

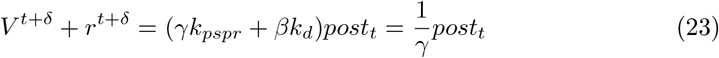

### 6.3 Synaptic Calcium model

The dynamic for dendrite potential is similar to the *ρ*

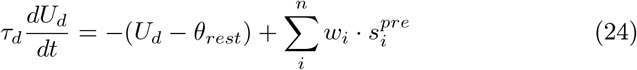

A pre-synpatic spike release glutamate to the synaptic cleft in the amount of *Q* called Quantal Content in literature [26]. Glutamate reuptake and decomposition act on a relatively long time scale.

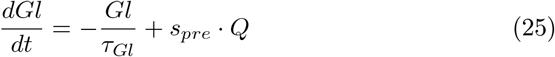

Glutamate can bind to NMDAR which is a calcium-permeable ion channel. NMDAR is blocked by magnesium that can be removed by depolarizing potential [21]. Its activation has a supra-linear voltage dependency modelled by two linear stages here. *U*_*d*_ exceeding the threshold *θ*_*high*_ entering high area. Calcium leaks out with a time constant *τ*_*Ca*_. [*x*]^+^ = *max*(*x*, 0) is for threshold activation.

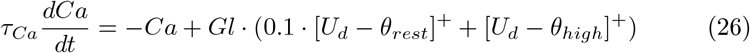

In order to model the hypothesis that moderate calcium concentration induces depression while high concentration leads to potentiation [38], we derive two separate processes for LTD/LTP with logarithm/linear dependency on calcium. LTD is proportional to *Ca* in logarithm and requires glutamate above threshold *θ*_*Gl*_.

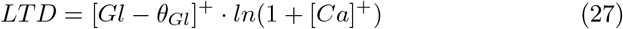

LTP is linear to *Ca* but with a higher threshold *θ*_*LT P*_.

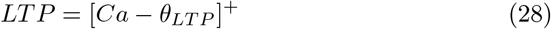

## 7 Pending extensive experiments

### 7.1 Maze task

A maze navigation task similar to [9] has been designed and implemented with PyTorch acceleration. We design state neurons like the place cell that maps to locations in the 2-D grid map. To explore TDSTDP as part of Actor-Critic as we discussed: we need to devise another neural network afferent also from state neuron and output from action neuron. This network is modulated by DA thus a three-factor plasticity rule is appreciated here.

### 7.2 Ramping DA release

Try to reproduce the DA positive ramping in [14] as the decay *γ* theoretically ensures a concave value function. It is also possible to adapt state neurons to Pavlov’s delayed reward mode, which should induce negative ramping instead. The cue should be represented by weakening state neurons.

